# Cell-Type-Specific Regulation of Cocaine Reward by the E2F3a Transcription Factor in Nucleus Accumbens

**DOI:** 10.1101/2024.07.08.602609

**Authors:** Freddyson J. Martínez-Rivera, Yun Young Yim, Arthur Godino, Angélica Minier-Toribio, Solange Tofani, Leanne M. Holt, Angélica Torres-Berrío, Rita Futamura, Caleb J. Browne, Tamara Markovic, Peter J. Hamilton, Rachael L. Neve, Eric J. Nestler

**Author notes:** Department of Neuroscience, University of Florida, Gainesville, Florida, USA. Co-first authors (Freddyson J. Martínez-Rivera, Yun Young Yim). **Co-correspondence** Freddyson J. Martinez-Rivera, PhD, Assistant Professor, Department of Neuroscience, McKnight Brain Institute, College of Medicine, University of Florida, Gainesville, Yun Young Yim, PhD, Instructor, Nestler Lab, Department of Neuroscience, Icahn School of Medicine at Mount Sinai, Eric J. Nestler, MD, PhD, Nash Family Professor of Neuroscience, Director, The Friedman Brain Institute, Dean for Academic and Scientific Affairs.

## Abstract

The development of drug addiction is characterized by molecular changes in brain reward regions that lead to the transition from recreational to compulsive drug use. These neurobiological processes in brain reward regions, such as the nucleus accumbens (NAc), are orchestrated in large part by transcriptional regulation. Our group recently identified the transcription factor E2F3a as a novel regulator of cocaine’s rewarding effects and gene expression regulation in the NAc of male mice. Despite this progress, no information is available about the role of E2F3a in regulating cocaine reward at the sex- and cell-specific levels. Here, we used male and female mice expressing Cre-recombinase in either D1- or D2-type medium spiny neurons (MSNs) combined with viral-mediated gene transfer to bidirectionally control levels of E2F3a in a cell-type-specific manner in the NAc during conditioned place preference (CPP) to cocaine. Our findings show that selective overexpression of E2F3a in D1-MSNs increased cocaine CPP in both male and female mice, whereas opposite effects were observed under knockdown conditions. In contrast, equivalent E2F3a manipulations in D2-MSNs had no significant effects. To further explore the role of E2F3a in sophisticated operant and motivated behaviors, we performed viral manipulations of all NAc neurons in combination with cocaine self-administration and behavioral economics procedures in rats and demonstrated that E2F3a regulates sensitivity aspects of cocaine seeking and taking. These results confirm E2F3a as a central substrate of cocaine reward and demonstrate that this effect is mediated in D1-MSNs, thereby providing increased knowledge of cocaine action at the transcriptional level.

## Introduction

Collective efforts in addiction research have shined light on the neurobiological mechanisms of a drug’s rewarding and reinforcing properties. Such outcomes have demonstrated the prominence of molecular changes, particularly including epigenetic and transcriptional processes, in mediating the lasting consequences of drug action [1]. Cocaine’s effects, in particular, have been associated strongly with key transcription factors within brain reward regions that shape gene expression patterns linked to the development of cocaine use disorders (CUD). Among these factors, we and other groups have identified ΔFOSB, CREB, EGR3, SRF, and more recently RXRA, ZFP189, and E2F3 [2–5].

E2F3 is involved in chromatin dynamics and gene/RNA regulation [6,7], with two isoforms (E2F3a and E2F3b) that have arisen as pivotal mediators of cocaine action in a region and isoform-dependent manner [8,9]. While E2F3a exerts its causal effects in promoting cocaine reward in the nucleus accumbens (NAc) [8], E2F3b displays similar effects in the prefrontal cortex (PFC) [9]. However, only E2F3a has been found to act as an upstream regulator of the *Fosb* gene and its protein product ΔFOSB [10], which is a canonical factor of cocaine-associated plasticity [11]. Consistent with this, overexpression of E2F3a in the NAc recapitulates the overall transcriptomic pattern induced in this brain region by chronic cocaine exposure [8]. In addition, genome-wide analyses implicate E2F3a acting in the NAc as one of the strongest upstream regulators of the transcriptomic and epigenetic actions of cocaine [10,12,13], effects that have recently been shown to recapitulate the molecular pathology in this brain region in humans suffering from CUD [14].

While most of these effects of E2F3a have been described in a non-cell-specific manner, our group found that the induction of *Fosb* gene expression in the NAc by E2F3a largely occurs in D1-medium spiny neurons (D1-MSNs) rather than in D2-MSNs [10]. Furthermore, all prior studies of E2F3a in cocaine action have been limited to male mice only. Thus, to expand the characterization of E2F3a at the sex- and cell-specific levels, we used viral-mediated gene transfer combined with cocaine conditioned-place preference (CPP) in transgenic male and female mice expressing Cre-recombinase in either D1- or D2-MSNs. To complement these approaches, we used cocaine self-administration and behavioral economic tasks as a first step to examine the role of E2F3a in operant and motivated behaviors in rats. Building upon existing data, our new findings support the importance of E2F3a as a “molecular switch” that is involved in the development of cocaine addiction.

## Methods

### Animals

#### Mice

Adult male and female transgenic mouse lines (D1-Cre: MGI:3836633, D2-Cre: MGI:3836635) were bred in-house on a C57BL/6J background. Mice (8-16 weeks old, 20-30 g), were grouped and housed on a 12 hr light-dark cycle (lights off at 19:00) with food and water ad libitum.

#### Rats

Adult male Sprague-Dawley rats (300-400 g; Charles River) were maintained on a 12 hr reverse light/dark cycle (lights on at 19:00) and restricted to 95% of their free-feeding body weight to facilitate self-administration behavior. Initially, rats were pair-housed until the self-administration procedures began, after which they were single housed.

All mouse and rat procedures were approved by the Institutional Animal Care and Use Committee (IACUC) of the Icahn School of Medicine at Mount Sinai and were in accordance with the guidelines of the Association for Assessment and Accreditation of Laboratory Animal Care (AAALAC) and the Society for Neuroscience.

### Virus

E2F3a overexpression (OE) and knockdown (KD) were performed using herpes simplex virus 1 (HSV)-based vectors (which are replication defective) [15–17] in a Cre-dependent manner. We used bicistronic p1005 HSV1 vectors (concatemer amplicon) expressing E2f3a (plus EGFP) or EGFP genes (control) for overexpression. In knockdown experiments, HSV vectors expressing microRNAs targeting E2f3a or LacZ (control) which also express EGFP were used. These experimental vectors expressed the genes under the cytomegalovirus (CMV) enhancer/promoter (for viral tropism) and were flanked with two inversed loxP sites (LS1L; Cre-dependent sites) that only express the transgene of interest driven by the IE4/5 promoter enforcing the infections of neuronal cells. Control constructs expressed EGFP in a Cre-dependent manner under the IE4/5 promoter, whereas in experimental viruses, EGFP induction was not Cre-dependent. The viruses were labeled as follows: Cre-dependent (mice); overexpression; HSV-LSL1-E2F3a-EGFP, HSV-LSL1-EGFP (control); knockdown: HSV-LSL1-mir-E2F3a-EGFP, HSV-LSL1-mir-LacZ-EGFP (control). These new vectors were thoroughly validated to confirm the cell-type-specific expression of their transgenes (Fig. S1). For non-conditional approaches, the following published viruses were used in rats: HSV-mir-E2F3a-EGFP, HSV-mir-LacZ-EGFP (control) [8]. Plasmid cloning and packaging into HSVs vectors were performed at the Gene Delivery Technology Core of Massachusetts General Hospital in Boston, Massachusetts, USA.

### Stereotaxic surgery and viral-mediated gene transfer

Mice or rats were intraperitoneally (i.p.) administrated with a mixture of ketamine (100 mg/kg) and xylazine (10 mg/kg) diluted in saline for general anesthesia. This was followed by head-fixation in a stereotaxic frame (Kopf instruments), where Hamilton syringes (33-gauge) were used to bilaterally infuse (1 ml; rate 0.1 ml/min) the viral vectors (±1×10^9^ IU/mL titers) in the NAc. The stereotaxic coordinates were as follows (relative to Bregma): mouse: +1.6 mm A/P, +1.4 mm M/L, −4.5 mm D/V, 10° angle; rat: +1.7 mm A/P, +2.3 mm M/L, −6.5 mm D/V, 10° angle. The virus was allowed to diffuse for 10 min before the needle removal. To allow maximal viral transduction, the CPP procedures started 4 days after the surgery in overexpression experiments, whereas in knockdown conditions 6 days were allowed before the CPP started (mice) or 2 days for the resumption of the self-administration task (rats) [3,8,9].

### Drug

Cocaine HCl (NIDA) was dissolved in 0.9% NaCl saline (ICU medical) for CPP (5 mg/kg and 7.5 mg/kg) [8] and self-administration experiments (0.8 mg/kg/infusion) [18,19].

### Conditioned-place preference

A three-chamber apparatus operated by Med Associates Inc. (MED-PC IV v4.2 software) was used for cocaine CPP as described previously [3,20,21]. Briefly, the protocol consisted of three phases; pre-test, conditioning, and post-test under semi-dark conditions. During the pre-test, mice were allowed to explore all three chambers for 20 min. Afterward, we unbiasedly counterbalanced the groups and randomly assigned the drug/saline sides. In the conditioning phase, mice were administrated (i.p.) with either saline (in the morning) or cocaine (in the afternoon) and confined to the corresponding CPP side for 30-min sessions over two days. On the following day (post-test), mice were retested and the preference score was recalculated as the difference between the time spent in the drug and saline sides

### Jugular vein catheterization (JVC) and cocaine self-administration

Rats were anesthetized with a mixture of ketamine (100 mg/kg) and xylazine (10 mg/kg) and implanted with chronic indwelling catheters (in-house made) into the right jugular vein [3,18,19]. An analgesic (ketoprofen; 5 mg/kg, Zoetis US) was subcutaneously administered along with topical antibiotic. Catheters were flushed daily with 0.1 ml heparinized saline (heparin; 30 U) containing ampicillin (5 mg/ml; Patterson) during the recovery phase (4-6 days) and with heparin for the remainder of the experiment. After the recovery period, rats were voluntarily trained for cocaine (0.8 mg/kg/infusion) self-administration under a fixed ratio one (FR1) during 3 hr sessions. Presses to the active lever (correct) resulted in an infusion (5 sec) that was followed by a 20 sec timeout period during which the levers were retracted and a cue light was illuminated. Presses on the inactive (incorrect) lever were recorded but had no consequences. Acquisition was considered established when >70% of the presses were correct. Behavioral recordings were obtained using Med Associates systems (operant boxes and Med PC interface).

### Behavioral economics threshold procedure

Upon acquisition of cocaine self-administration, rats received a pre-test session of behavioral economics to counterbalance and assign the groups based on their pre-existent motivational thresholds prior to viral-mediated gene transfer. Following stereotaxic surgery (viral infusion), rats received two behavioral economics sessions, where each session was preceded by an FR1 session on different days. The incorporation of FR1 days helped to reduce any extinction learning and to reestablish the acquired level of seeking behavior affected by the demand of the behavioral economics task. Behavioral economics was performed as previously described by our group with minor modifications [3,18,19]. Briefly, the task consisted of 110 min sessions with 11 descending doses of cocaine (259.88, 146.14, 82.18, 46.22, 26.00, 14.62, 8.23, 4.61, 2.61, 1.46, 0.83 µg/infusion) in which each dose was available for self-administration during 10 min in an FR1 ratio with no timeouts and levers extended. Lever presses and cocaine intake were represented and plotted as a function of dose/price, where the price was defined as the number of responses required to obtain 1 mg of cocaine.

For the resulting demand curves, we performed the curve fitting method using the following mathematical equation: log_10_(*Q*) = log_10_(*Q*_0_) + *kx*(*e*^−α*Q*^_0_C−1). Here, Q is intake and C is the varying cost of cocaine [3,18,19]. The *k* value, which is the number of estimated parameters, was set to 2 by default but was determined to be 2.7 by fitting all individual curves with increments of 0.1. The curve fitting was performed using a custom-made R package that applies the stats::nls function, where the goodness of fit was obtained by calculating a pseudo-R2 coefficient, which is the square of the correlation between the observed and predicted log_10_(Q) values and the Akaike Information Criterion (AIC) for each fit [3].

In the initial bins where the high doses are available, the intake at the minimal imposed price can be derived and defined as Q_0_. As the cocaine doses progressively decrease, the cost/price increases to the level where the animals will no longer maintain their responses, leading to a decrease in cocaine intake. This is defined by the maximal price paid, denoted as P_max_ to maintain Qo responding. Responses during the first dose (bin) that reflect the loading/priming period, were not included in the formal analysis. The Q_max_ and O_max_ parameters represent the intake and active lever presses at the P_max_, respectively, and these parameters were calculated for each rat [3,18,19]. As a measure of how sensitive the rats are to changes in the price demand, the behavioral elasticity (α) was calculated. A higher α indicates a quicker drop in responses will be observed (elastic behavior), whereas a lower α defines inelastic behavior. Curve fitting analysis and R code are available upon request [3].

### Tissue collection and placement verification

After the CPP experiments (24-48 hr), mice were cervically dislocated, and their brains were immediately extracted and placed in chilled PBS. Subsequent coronal (1 mm) brain sections were obtained and viral (EGFP) expression and spread in the NAc were confirmed using a fluorescent microscope (Leica SP8). Bilateral punches (12-gauge) of the NAc were collected and stored at −80°C for any additional analysis. A separate subset of mice was used to validate viral overexpression or knockdown using RT-PCR (Fig. S1).

In the self-administration experiment in rats, where the non-conditional virus had been previously validated [8,9], brain fixation and sectioning were performed to verify viral expression and spread. After the last behavioral economics session (24-48 hr), rats were injected with sodium pentobarbital (fatal plus: 450 mg/kg i.p.) and were perfused transcardially with cold PBS followed by cold freshly prepared 4% paraformaldehyde (PFA) in potassium phosphate buffer (PBS). Brains were then extracted and stored in 4% PFA (24-48 hr) before being transferred into a 4% PFA + 30% sucrose solution for cryoprotection for at least 24-48 hr. Brains were then sectioned (40 µm) using a cryostat (Leica), mounted in supercharged slides (Fisher Scientific), and observed under a fluorescent microscope (Zeiss Imager. M1). A separate subset of mice was also perfused to visualize viral placement (Fig. 1C).

**Figure 1.**
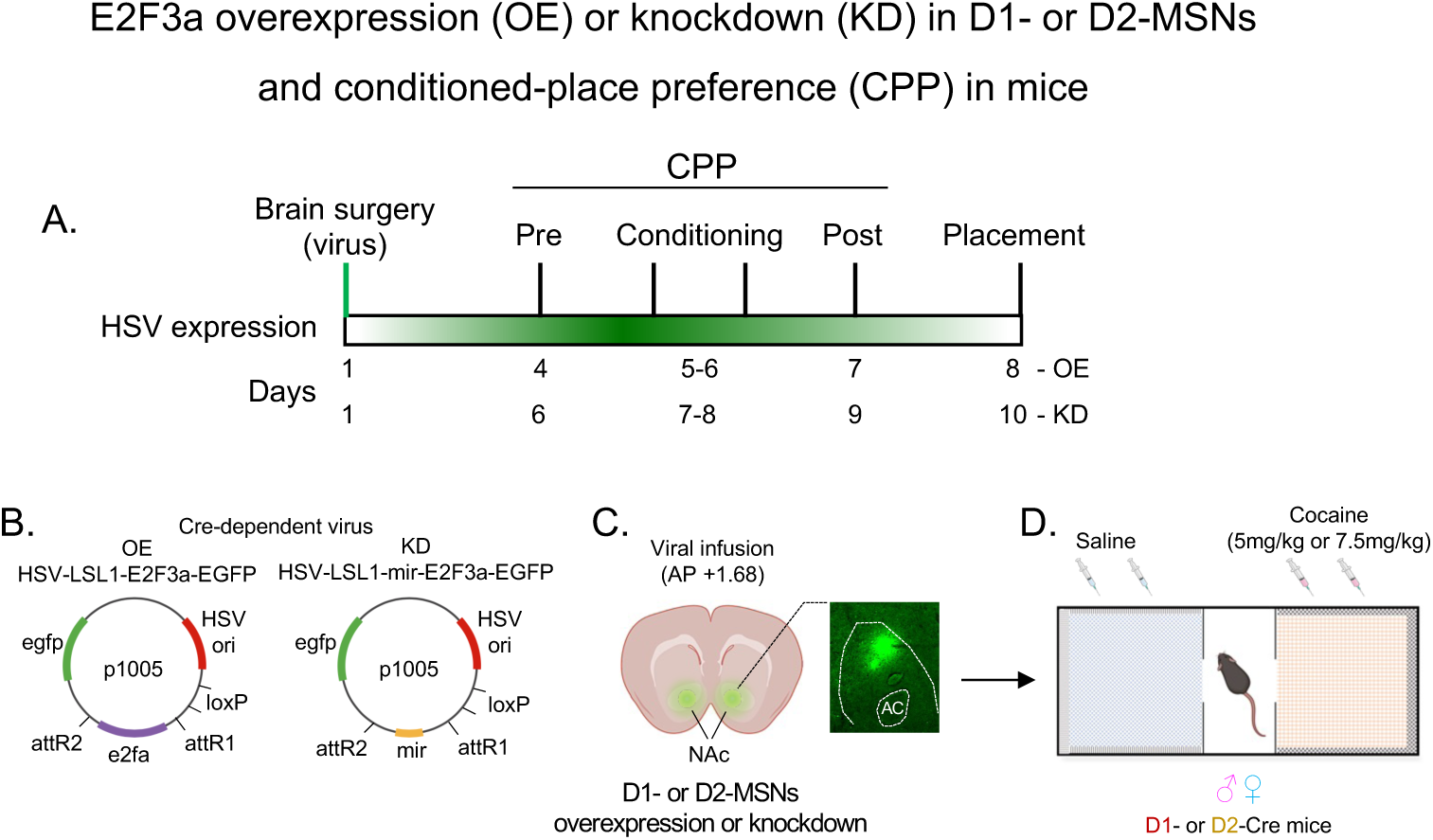
Experimental design of viral-mediated gene transfer and CPP. **A.** Timeline of the stereotaxic surgeries and viral infusion into the NAc and the CPP phases under E2F3a overexpression or knockdown conditions. **B-C.** Schematics illustrating the Cre-dependent vectors (backbones) for overexpressing or knocking down E2F3a to be infused in the NAc (with a representative placement schematic and micrograph). **D.** The CPP paradigm in male and female mice expressing Cre-recombinase in either D1- or D2-MSNs and the doses utilized for overexpression (5 mg/kg) and knockdown (7.5 mg/kg) experiments. AC = Anterior commissure.

### Real-time polymerase chain reaction (RT-PCR)

Frozen NAc punches were prepared for RNA isolation using the Zymo Direct-zol MiniPrep kit according to the manufacturer’s instructions. Samples were then quantified using a NAnoDrop 2000 (Thermofisher). Total RNA (150 ng) was converted to cDNA using the Bio-Rad iScript cDNA Synthesis Kit. qPCR reactions were performed in triplicates using SYBR green (Applied Biosystems) and Applied Biosystems 7900HT RT-PCR machine with the following parameters: 2 min at 95 °C; 40 cycles of 95 °C for 15 sec, 58 °C for 30 sec, and 72 °C for 33 sec. Data analysis was performed using the 2^-ΔΔCt^ method [22] from threshold cycle (Ct) values comparing GAPDH (F: AGG TCGGTGTGAACGGATTTG / R:TGTAGACCATGTAGTTGAGGTCA IDT) controls vs E2F3a (F: CTACACCACGCCACAAGGAC / R: CTCCGTAGTGCAGCTCTTCC IDT) OE or KD samples (Fig. S1C-D) [8].

### Statistical analyses

Statistical analyses, including two-way repeated measures (RM) ANOVAs, followed by Sidak’s post-hoc tests, were performed for CPP and self-administration (behavioral economics) results. Student’s two-tailed t-tests were also performed for behavioral economics (Welch correction) and RT-PCR results. Data are presented as mean ± standard error (SEM) and statistical significance was established as *p<0.05 (Prism 9.3.1 software). The CPP score was defined as: time spent on the drug side - time spent on the saline side.

## Results

### E2f3a drives cocaine reward via D1-MSNs of the NAc in male and female mice

To test whether E2F3a exerts its effects on cocaine reward in a sex- and cell-specific manner, we performed cocaine CPP in male and female mice expressing Cre-recombinase in either D1- or D2-MSNs. This was combined with viral-mediated gene transfer using Cre-dependent HSV vectors to induce E2F3a overexpression (OE) or knockdown (KD) in the NAc (Figs. 1A-D, S1A-D). Mice were then pre-tested to counterbalance the groups and randomly assigned in an unbiased fashion the cocaine/saline-paired sides based on animals’ baseline preferences for the CPP compartments. Conditioning was conducted over the subsequent two days with saline (in the morning) and cocaine (in the afternoon), followed by a post-test the next day. As an index of conditioning, the time spent in the cocaine- and saline-paired sides during both the pre- and post-tests were calculated to determine a place preference score.

In overexpression conditions, where E2F3a was hypothesized to facilitate cocaine CPP, a lower dose (5 mg/kg) – which is normally subthreshold for cocaine conditioning – was administered. When pre- and post-tests were compared within groups, only mice overexpressing E2F3a in D1-MSNs showed an increase in cocaine place preference, an effect that was apparent in male and female mice (time: males: F_(1,30)_ = 4.19, p = 0.04, Sidak post-hoc; EGFP: p = 0.99, E2F3a: p = 0.02; females: F_(1,39)_ = 4.37, p = 0.04, Sidak post-hoc (pre vs. post), EGFP: p = 0.83, E2F3a: p = 0.05; Fig. 2A-B). Notable trends between group x time interaction were also observed (males: F_(1,30)_ = 3.61, p = 0.06; females: F_(1,39)_ = 1.88, p = 0.1). Between group (virus) comparisons revealed elevated (but not significant) post-test values under OE conditions (post: males: EGFP: 35.4 ± 64.5; E2F3a: 218.6 ± 49.9; F_(1,30)_ = 1.46, p = 0.23, Sidak post-hoc, p = 0.10; females: EGFP: 34.6 ± 31.1; E2F3a: 158.1 ± 44.3; F_(1,39)_ = 0.90, p = 0.34, Sidak post-hoc, p = 0.20). While these effects were more robust in males, no sex differences were observed when directly comparing both sexes, virus, and time (pre and post-tests) variables (sex: F_(1,69)_ = 0.66, p = 0.41).

**Figure 2.**
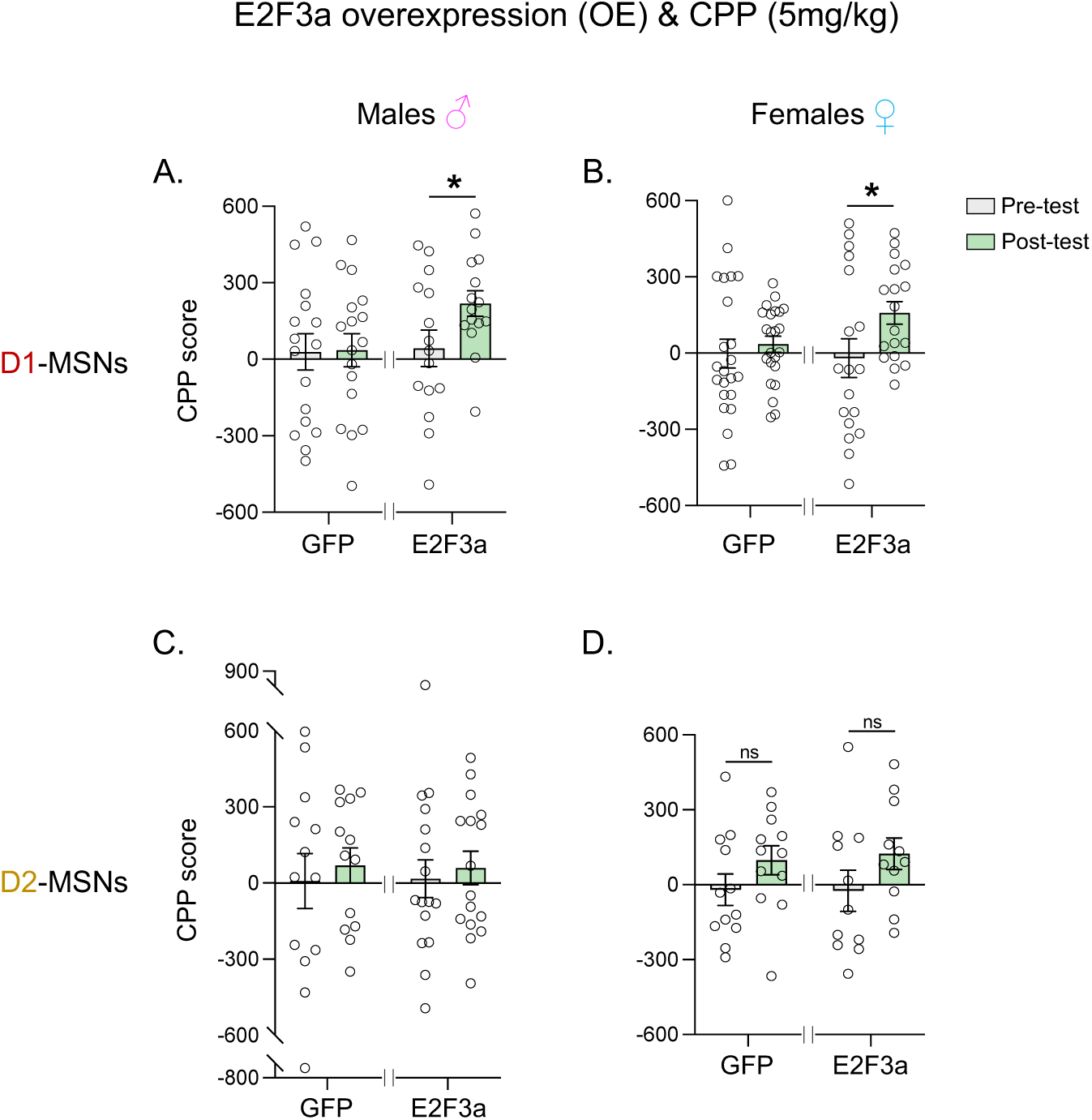
E2F3a overexpression in D1-MSNs but not D2-MSNs facilitates cocaine CPP in mice. **A-B.** Following the pre-test and conditioning phases of cocaine CPP at a subthreshold dose (5 mg/kg), overexpression (OE) of E2F3a in D1-MSNs of the NAc facilitated cocaine CPP (post-test). This effect increased the time spent on the cocaine-paired side during the post-test in both male (EGFP n = 17; E2F3a/OE n = 15) and female (EGFP n = 23; E2F3a/OE n = 18) mice. **C-D.** Overexpressing E2F3a in D2-MSNs had no significant effect as observed by the lack of CPP for a subthreshold dose in either males (EGFP n = 13; E2F3a/OE n = 16) or females (EGFP n = 12; E2F3a/OE n = 11) mice. All data are shown as mean ± SEM. \**p* < 0.05, ns = non-significant.

In contrast, E2F3a overexpression in D2-MSNs did not promote cocaine CPP in either sex (Fig. 2C-D). Although a small CPP increase was observed under GFP and E2F3a conditions in female mice, post-hoc analyses revealed no significant differences between pre- and post-test scores in either sex with either virus (time: males: F_(1,27)_ = 0.83, p = 0.36, Sidak post-hoc (pre vs. post), EGFP: p = 0.72, E2F3a: p = 0.82; females: F_(1,21)_ = 7.37, p = 0.01; Sidak post-hoc (pre vs. post), EGFP: p = 0.2, E2F3a: p = 0.1. Virus (group): males: F_(1,27)_ < 0.001, p = 0.99, Sidak post-hoc, post-test (EGFP vs. E2F3a): p = 0.9; females: F_(1,21)_ = 0.01, p = 0.89, Sidak post-hoc, post-test (EGFP vs. E2F3a): p = 0.9; Fig. 2C-D). Consistent with this no sex differences were observed when directly comparing both sexes, virus, and time (pre- and post-tests) variables (sex: F_(1,48)_ < 0.01, p = 0.92). Together, these results demonstrate a clear cell-type difference in the actions of E2F3a and establish that E2F3a acting in D1-MSNs is sufficient to promote cocaine reward in a non-sex-dependent manner.

To test the converse, whether E2F3a is also necessary for the expression of cocaine CPP, a higher dose of cocaine was used (7.5 mg/kg) under knockdown conditions. Here, viral-mediated gene transfer was performed to knockdown E2F3a in a sex- and cell-specific manner. When E2F3a was silenced in D1-MSNs, pre- and post-test comparisons revealed that only control (mir-lacZ) mice developed cocaine CPP, an effect seen in both male and female mice (time: males: F_(1,16)_ = 4.15, p = 0.05, Sidak post-hoc (pre vs. post); mir-lacZ: p = 0.03, mir-E2F3a: p = 0.99; females: F_(1,20)_ = 2.04, p = 0.16, Sidak post-hoc (pre vs. post); mir-lacZ: p = 0.04, mir-E2F3a: p = 0.82; Fig. 3A-B). Additional group comparisons also highlight attenuated CPP under KD conditions in both sexes (group (post-test): males: mir-lacZ: 280.3 ± 71.1; mir-E2F3a: 4.1 ± 86.9; F_(1,16)_ = 1.89, p = 0.19, Sidak post-hoc (lacZ vs. E2F3a), p = 0.06; females: mir-lacZ: 118.5 ± 52.7; mir-E2F3a: −52.5 ± 59.9; F_(1,20)_ = 1.61, p = 0.22, Sidak post-hoc (lacZ vs. E2F3a), p = 0.07. Such findings were accompanied by notable interactions between group x time for both sexes (males: F_(1,16)_ = 4.06, p = 0.06; females: F_(1,20)_ = 4.72, p = 0.04). No sex differences were observed when directly comparing both sexes, virus, and time (pre and post-tests) variables (sex: F_(1,36)_ = 1.66, p = 0.27).

**Figure 3.**
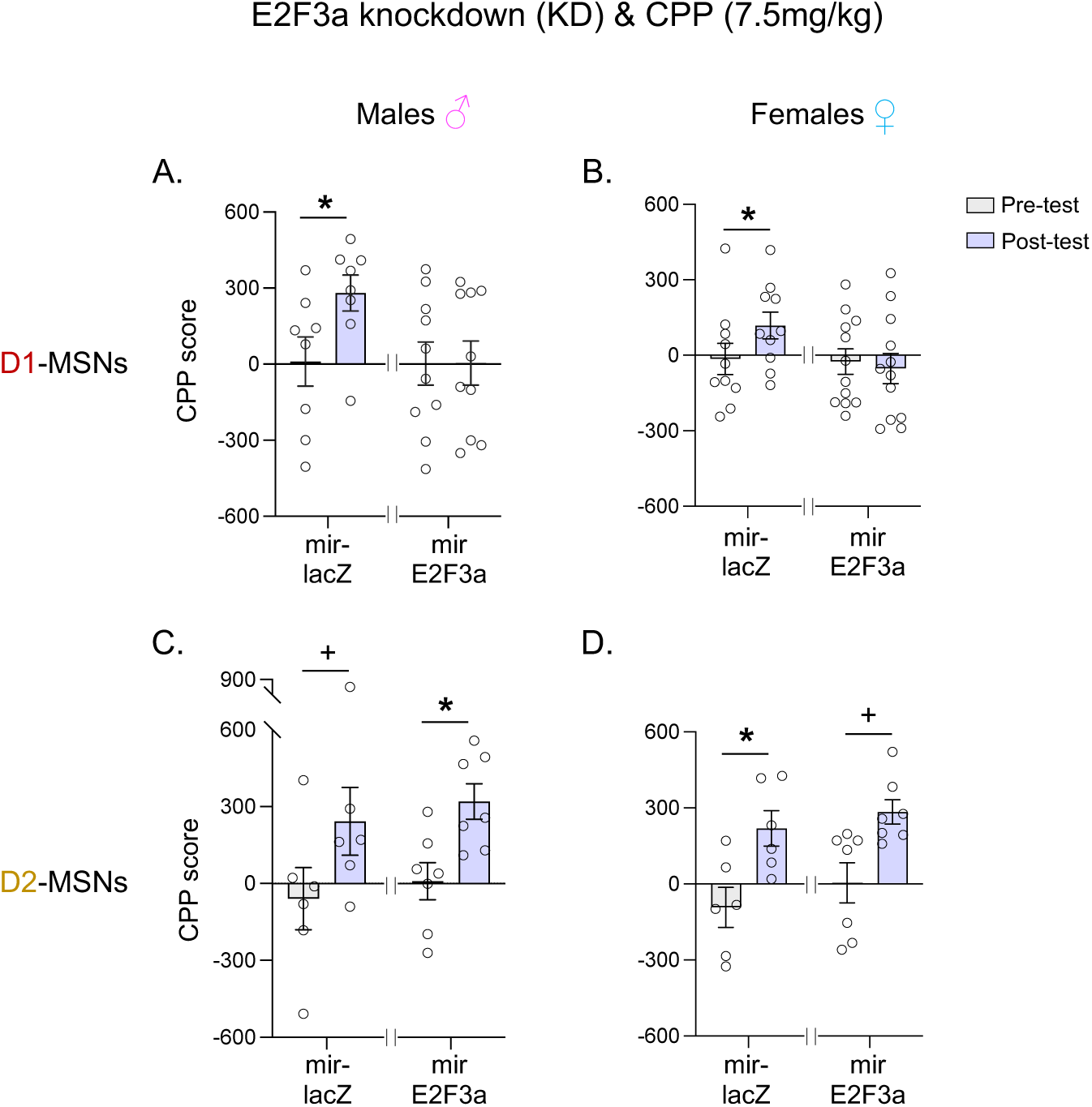
E2F3a knockdown in D1-MSNs but not in D2-MSNs prevents cocaine CPP in mice. **A-B.** Following the pre-test and conditioning phases of cocaine CPP at a higher dose (7.5 mg/kg), E2F3a knockdown (KD) in D1-MSNs of the NAc prevented cocaine CPP during the post-test. This was evidenced by a reduced time spent on the drug side, an effect observed for both male (lacZ n = 8; mir/KD n = 10) and female (lacZ n = 10; mir/KD n = 12) mice. **C-D.** Suppressing E2F3a in D2-MSNs had no significant effect as observed by the expected cocaine CPP for higher drug dose in both sexes (males: lacZ n = 6; mir/KD n = 7; females: lacZ n = 6; mir/KD n = 7). All data are shown as mean ± SEM. \**p* < 0.05, +*p ≤* 0.1.

On the other hand, such outcomes were not observed when E2F3a expression was knocked down in D2-MSNs: cocaine CPP was unchanged in both male and female mice (time: males: F_(1,11)_ = 12.61, p < 0.01, Sidak post-hoc (pre- vs. post-tests), mir-lacZ: p = 0.07, mir-E2F3a: p = 0.04; females: F_(1,11)_ = 13.49, p < 0.01, Sidak post-hoc (pre- vs. post-tests), mir-lacZ: p = 0.04, mir-E2F3a: p = 0.05. Fig. 3C-D). Consonant with this, no evident differences by virus (lacZ vs. E2F3a) were observed in both sexes (males: group: F_(1,11)_ = 0.43, p = 0.52; interaction: F_(1,11)_ = 0.00, p = 0.96; females: group: F_(1,11)_ = 2.04, p = 0.18; interaction: F_(1,11)_ = 0.03, p = 0.84) or when directly comparing both sexes, virus, and time (pre and post-tests) variables (sex: F_(1,22)_ = 0.15, p = 0.70). These findings corroborate the cell-specific actions of E2F3a and establish that this transcription factor in D1-MSNs of the NAc is also required for normal rewarding responses to cocaine in both male and female mice.

### E2f3a controls cocaine consumption and motivation

While the CPP paradigm is used as a readout for associative learning and as an indirect measure of the rewarding (or aversive) properties of a drug, it is limited in measuring motivation, drug sensitivity, effort, and volitional aspects of drug seeking and taking (i.e., intake). To address the role of E2F3a in these far more complex behaviors, we conducted cocaine self-administration and a behavioral economics threshold procedure in rats. These studies were combined with viral-mediated knockdown of E2F3a in all NAc neurons to test whether this transcription factor is necessary for aspects of cocaine reinforcement (Fig. 4A-B). Rats were used for these experiments because they exhibit more robust self-administration behavior.

**Figure 4.**
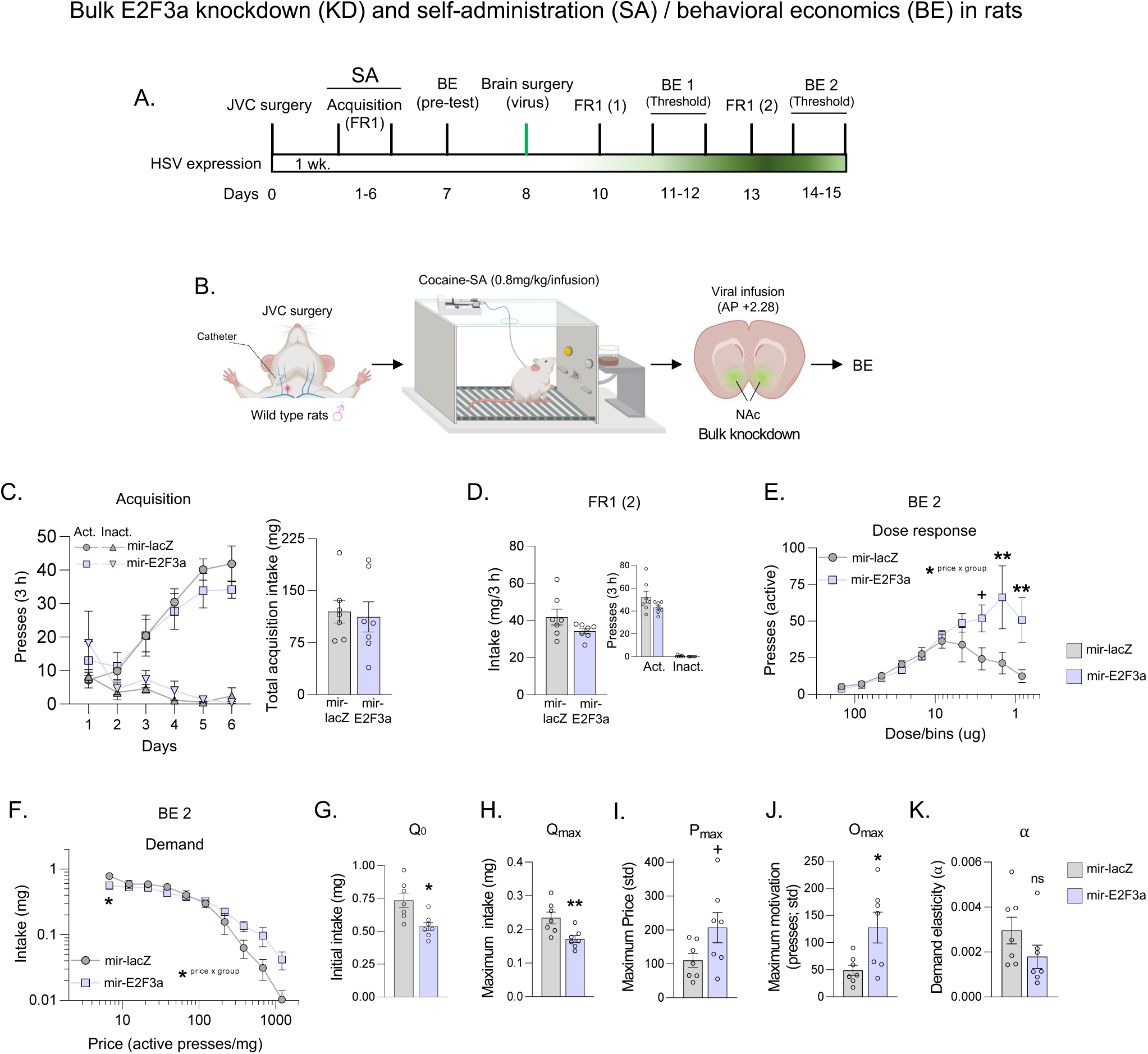
Knocking down E2F3a expression in the rat NAc alters motivation and intake features of cocaine self-administration. **A-B.** Schematics showing the timeline for jugular vein catheterization (JVC), cocaine self-administration, stereotaxic surgeries (virus infusion; knockdown; KD), and behavioral economics (BE) threshold procedures in rats. **C.** Acquisition of cocaine self-administration as measured by elevated active lever pressing and intake as well as reduced inactive lever pressing. **D.** No changes in cocaine self-administration were observed during the first FR1 session. **E.** Dose-response curve in the BE threshold procedure showed increased response for low doses of cocaine under KD conditions. **F.** Bidirectional demand curve shows decreased cocaine intake at a lower price (higher dose) and increased intake at a higher price (lower dose). **G-K.** These panels depict the specifics of BE performance measured by various parameters, including intake and motivation metrics/derivatives (plotted in this order), such as consumption at low effort (Q_0_), consumption at maximum effort (Q_max_), maximum price paid (P_max_), maximum lever presses (O_max_), and demand elasticity (α). A general decrease in consumption (Q’s) and increased motivation (P_max_, and O_max_) were seen under KD conditions. Such results were accompanied by less elasticity (α) under KD conditions as well. mir-lacZ n = 7; mir-KD n = 8. All data are shown as mean ± SEM. \*\**p ≤* 0.01, \**p* < 0.05, +*p ≤* 0.1.

Upon the acquisition of cocaine self-administration (Fig. 4C), rats were counterbalanced and assigned to either control or experimental groups (virus) based on their initial behavioral profiles (behavioral economics pre-test/baseline), with no group differences in any of the measured parameters (Fig. S2A-H). This was followed by viral infusions (Fig. 4A-B), recovery time, sequential FR1 sessions 1 and 2, and behavioral economics sessions 1 and 2 (BE1 and BE2) (Fig. S2A). Major differences were observed during BE2 sessions, where maximal viral expression was achieved, although results during BE1 trended toward anticipating these differences (Fig. S2I-P). During the behavioral economics task, rats self-administered the drug at an increasing cost (“price”) due to decreasing the dose as the session progressed, thereby increasing their responses (presses) to the limit of drug seeking. As a result, dose-response and demand curves (and derivatives) were obtained.

Initially, during FR1 (session 2), no significant differences were captured in cocaine-seeking and -taking metrics under fixed contingencies (intake: t_(7)_ = 1.67, p = 0.13; active: t_(7)_ = 1.67, p = 0.13; inactive: t_(6)_ = 1.92, p = 0.11. Fig. 4D). However, during the subsequent behavioral economics session (BE2), when viral expression had peaked, dose-response analysis revealed an up/right shift/pattern as the session progressed. These results indicated higher responses at lower drug doses when E2F3a KD was compared with lacZ controls (group/virus; F_(1,12)_ = 3.51 p = 0.08, Sidak post-hoc; last three bins: p = 0.1 and p’s < 0.01. Fig. 4E). Interestingly, at a higher dose (first bin), fewer but not significant responses were observed under knockdown conditions (mir-lacZ: 5.35 ± 0.48; mir-E2F3a: 3.85 ± 0.48). As expected, both groups responded to the imposed doses across the session (dose effect: F_(9, 108)_ = 10.47, p < 0.01). In addition, these analyses showed a strong interaction between group and dose (F_(9, 108)_ = 4.07, p < 0.01), suggesting that deficient E2F3a expression in NAc cells alters sensitivity to cocaine (the higher the dose, the lower the responses, and vice versa). These findings are consistent with the CPP results, which showed an elevated place preference with a lower drug dose under overexpression conditions, and the opposite with a higher dose under knockdown conditions.

We next examined the demand curves (Fig. 4F), with cocaine intake as a function of price, where independent measures of motivation, consumption, and elasticity were derived using mathematical extrapolations of economic indicators (Fig. 4G-K). No significant differences between groups were noted (F_(1, 12)_ = 0.29, p = 0.5); however, post-hoc analyses revealed that silencing E2F3a in the NAc reduced intake at higher drug doses when the price was low (Sidak; first bin p < 0.01. Fig. 4F). Furthermore, a significant effect was observed as the price changed (F_(9, 108)_ = 71.10, p < 0.01), along with a strong interaction between groups and price (F_(9, 108)_ = 2.95, p < 0.01). Such results are consistent with the dose-response findings, showing that knocking down E2F3a in this brain region reduced intake at higher drug doses when the price was low, but increased the effort to consume more cocaine as the price increased, when the drug availability was limited (Sidak; first bin p < 0.01. Fig. 4F).

Derived behavioral economics parameters additionally showed that cocaine intake at both low (Q_0;_ t_(9)_ = 3.12, p = 0.01) and maximum (Q_max;_ t_(9)_ = 3.07, p = 0.01) effort was decreased upon E2F3a knockdown in the NAc (Fig. 4G-H). In contrast, a trending increase in motivation was observed under knockdown conditions as measured by P_max_ values (t_(8)_ = 1.97, p = 0.08. Fig. 4I), which reflects the maximum price the animals were willing to pay. Along with this finding, the maximum behavioral output (O_max_) spiked (t_(7)_ = 2.64, p = 0.03. Fig. 4J), supporting the idea that impairing E2F3a expression may prime the animal’s motivation to be more sensitive to cocaine, bidirectionally impacting cocaine seeking and taking behaviors as the available dose varies. Along the same line, an inelastic-like behavior (reduced trend of α) accompanied these results (mir-lacZ: 0.002 ± 0.0005; mir-E2F3a: 0.001 ± 0.0005; t_(11)_ = 1.48, p = 0.1. Fig. 4K), further highlighting an elevated effort to maintain the desired dose of cocaine as the price increases under knockdown conditions, an effect on intake that was not observed in FR1 sessions. Together, these results indicate a dynamic role of E2F3a in modulating dose/demand fluctuations, thereby fine-tuning an animal’s hedonic set point for cocaine consumption.

## Discussion

Building upon previous behavioral and transcriptomic studies that highlight E2F3a acting in the NAc as an important contributor to cocaine effects [8,10], we utilized cocaine CPP in male and female mice expressing Cre-recombinase in either D1- or D2-MSNs. This approach was combined with viral-mediated gene transfer in the NAc to overexpress or knockdown E2F3a in a sex- and cell-specific manner. We found that E2F3a selectively drives cocaine CPP via D1-MSNs in both sexes, where it is both necessary and sufficient for normal rewarding responses to cocaine. Our new CPP data align with an earlier observation that E2f3a induces ΔFOSB selectively in D1-MSNs of the NAc [10]. To further understand the role of this transcription factor in addiction-relevant behaviors, we performed a behavioral economics threshold procedure in rats while knocking down E2F3a in all NAc neurons. Together, our observations establish that E2F3a in the NAc strongly contributes to the reinforcing and motivational aspects of cocaine action.

While prior studies of E2F3a and cocaine focused on male animals only [8,10], results of the present investigation demonstrate an equivalent role for this transcription factor – acting selectively in D1-MSNs – in promoting rewarding responses to cocaine in male and female mice. This equivalency is striking given a growing literature that reveals prominent sex differences both in behavioral responses to cocaine and in the associated transcriptomic regulation in the NAc and other brain reward regions [23–25]. Still, similar actions of transcription factors in males and females is not without precedence, with ΔFOSB and RXRA as just two examples also exerting similar effects in the two sexes [3,26]. We recently used CUT&RUN sequencing (cleavage under targets and release using nuclease), which offers several advantages over ChIP-sequencing [27], to demonstrate highly overlapping direct genomic targets for ΔFOSB in the male and female NAc [26]. While it is unfortunately not feasible currently to apply CUT&RUN to E2F3a due to the lack of availability of suitable antibodies, such an approach might shed light on the precise mechanisms by which E2F3a acting in D1-MSNs induces similar effects on cocaine-regulated behaviors in males and females.

The conditioning and post-conditioning phases of CPP involve learning and memory components that are required to express the learned association [28]. Thus, performing manipulations prior to the conditioning phase can potentially impact both associative and expression (post-conditioning) phases, thereby masking the specific role of the targeted factor. Since our current and previous studies utilized viral manipulations before the commencement of the conditioning phase, addressing this issue requires manipulations both before and after this phase. The use of operant tasks (e.g., self-administration) would further help dissect these questions. For instance, differences in acquisition and seeking (expression) tests have been observed after virally manipulating other transcription factors relevant to cocaine-seeking behaviors at different time points [29].

To expand upon the CPP results, we tested the motivation to seek and take cocaine despite the cost, a common feature of addiction-like behaviors. We found that knocking down E2F3a in all NAc neurons affected motivation for and intake of cocaine in self-administration procedures as the dose and price of the drug varied. These behavioral economics data suggest that E2F3a might primarily affect an animal’s sensitivity to cocaine, resulting in complex effects in various aspects of self-administration behavior. Although more experiments are needed to determine the exact mechanisms underlying E2F3a action, one possibility is that some of its effects are mediated via induction of ΔFOSB. ΔFOSB has been shown to regulate numerous downstream genomic targets [26,30,31], many of which orchestrate synaptic plasticity and glutamatergic neurotransmission that are implicated in cocaine action [32,33]. Interestingly, studies in cancer research have suggested that E2F3a regulates sensitivity to chemotherapeutic drugs via epigenetic and apoptotic mechanisms [34–38], effects that could be extended to cocaine or other drugs of abuse.

Consistent with these findings with cocaine, studies in other addiction models have also highlighted the E2F family of transcription factors as playing an important role in substance use disorders (SUDs). For example, transcriptomic approaches combined with behavioral paradigms in mice linked several E2F isoforms with fentanyl and morphine abstinence [39], as well as with the risk of opioid use (e.g., heroin) [40]. Similar associations have been observed following exposure to other drugs such as methamphetamine, alcohol, and nicotine [41–45]. Our study shows that investigating the role of E2F isoforms, including E2F3a, in a sex-, cell-, and drug-dependent manner will be crucial for understanding their contributions across the spectrum of SUDs. Indeed, we showed previously that the E2F3b isoform had no discernable effect in the NAc, but potentiated cocaine CPP in the prefrontal cortex, whereas E2F3a had no discernable effect in this latter brain region [9]. Continued behavioral, transcriptomic, and epigenetic characterization of this transcription factor family will provide an increasingly robust neurobiological foundation for future interventions.

## Acknowledgments

This work was supported by NIH Grants: T32DA007135-33 & K01DA054306 to FJMR, R01DA007359 & P01DA047233 to EJN, T32DA053558 to LMH, and F99NS129172 to AMT. The authors want to thank Joseph Landry and James Callens (Yasmin Hurd Lab; self-administration procedures), as well as Giselle Rojas, Corrine Azizian, Kyra Schmidt, Nathalia Pulido, Katherine Beach, Stephen Pirpinias, Clementine Blaschke, Kinneret Rosen, and Ezekiell Mouzon (Nestler Lab) for their technical support (e.g., transgenic mouse breeding and genotyping) and their scientific feedback. We also thank the Gene Technology Core at Massachusetts General Hospital for HSV1 vectors. The authors declare no conflict of interest.

## Figure Legends for Supplementary Figures

**Figure S1.**
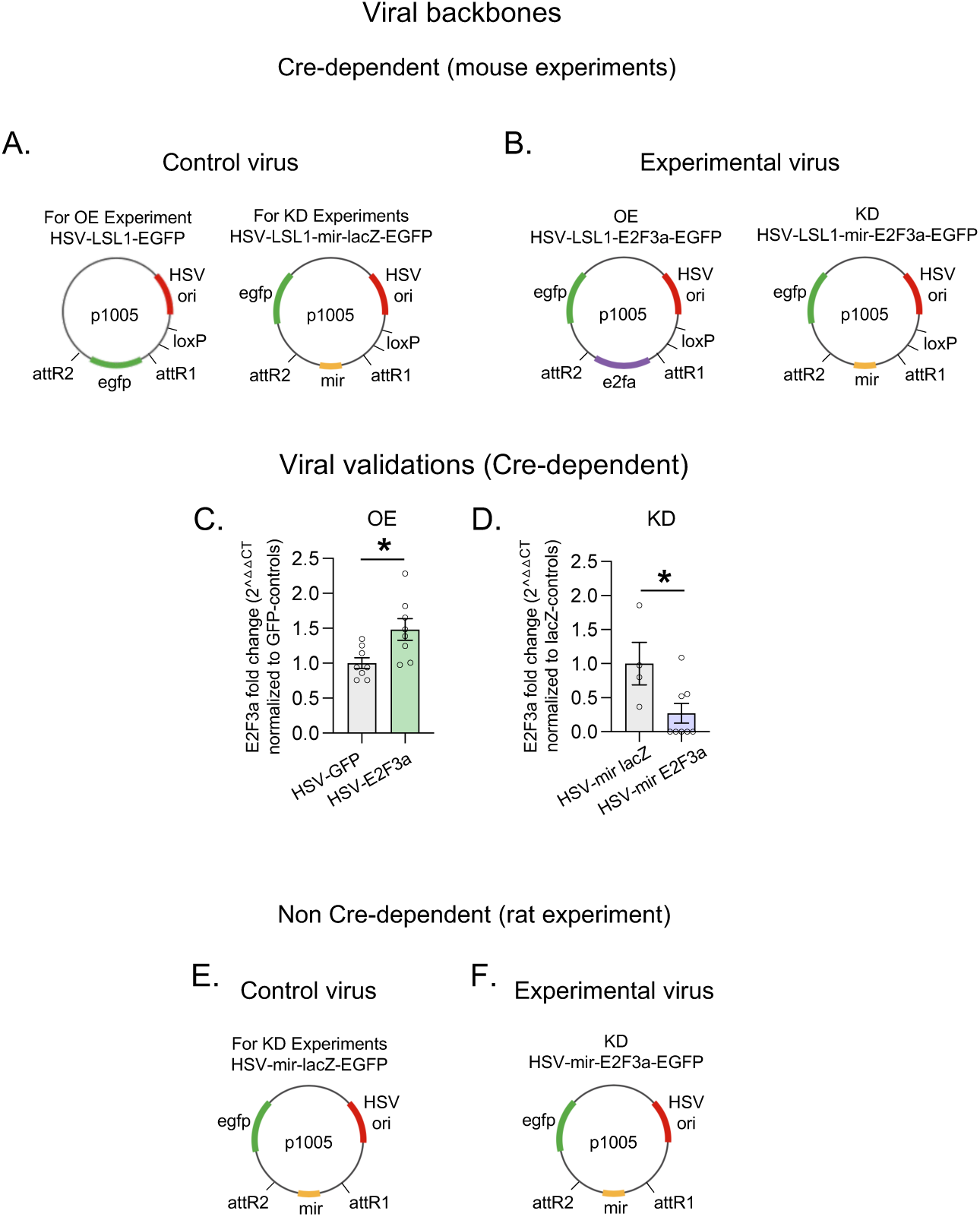
Validation of Cre-dependent HSV vectors. **A-B.** Schematics of the Cre-dependent vectors (backbones) overexpressing (OE) or knocking down (KD) E2F3a and control vectors. **C-D.** RT-PCR of *E2f3a* mRNA in the NAc showing fold change increase or decrease under OE or KD conditions, respectively. **E-F.** Schematics of the non-Cre-dependent vectors (backbones) for knocking down (KD) E2F3a and control vector. OE: D1-Cre females: EGFP n = 8; E2F3a/OE n = 8; t_(14)_ = 2.7, p = 0.01. KD: D1-Cre females: EGFP n = 4; E2F3a/KD n = 8; t_(10)_ = 2.45, p = 0.03. All data are shown as mean ± SEM. Unpaired t-test, \**p* < 0.05.

**Figure S2.**
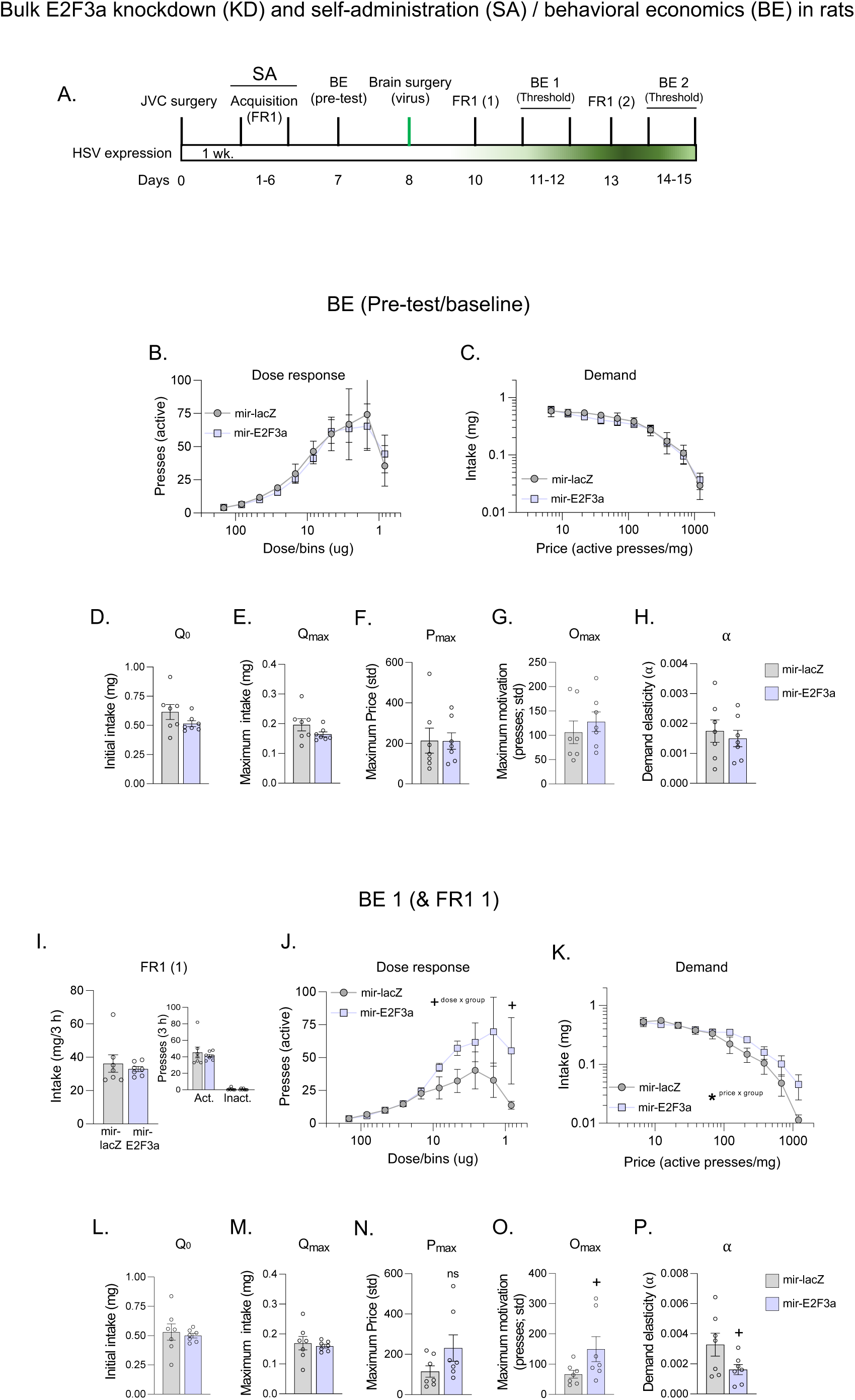
Initial behavioral economics matching/screenings. **A.** Schematics showing the timeline for jugular vein catheterization (JVC), cocaine self-administration, stereotaxic surgeries (virus infusion; knockdown; KD), and behavioral economics (BE) threshold procedures in rats. **B-H.** Matching/counterbalance of the assigned control and experimental groups, showing no basal differences in dose-response and demand curves and their derivatives (pre-BE). **I.** No changes in cocaine self-administration were observed during the first FR1 session of BE session 1 (BE1). **J.** Dose-response curve of BE1 showed an increasing trend in responses for low doses of cocaine under KD conditions accompanied by a trending interaction (dose x group; F_(9, 108)_ = 1.80, p = 0.07; dose: F_(9, 108)_ = 8.99, p < 0.01; group/virus: F_(1, 12)_ = 2.46, p > 0.05). **K.** Demand curve showing a higher intake as the price increases, with a significant interaction (price x group; dose x group; F_(9, 108)_ = 2.18, p = 0.02; price: F_(9, 108)_ = 76.99, p < 0.01; group/virus: F_(1, 12)_ = 0.49, p > 0.05). **L-P.** Panels depicting the specifics of BE performance as measured by different parameters, including intake and motivation metrics (plotted in this order) such as consumption at low effort (Q_0_), consumption at maximum effort (Q_max_), maximum price paid (P_max_), maximum lever presses (O_max_), and demand elasticity (α). Trends for increasing motivation (P_max_: t_(8)_ = 1.63, p = 0.14; and O_max_: t_(7)_ = 1.91, p = 0.09) and less elasticity (t_(8)_ = 1.98, p = 0.08) were observed under KD conditions. mir-lacZ n = 7; mir-KD n = 8. All data are shown as mean ± SEM. \**p* < 0.05, +*p ≤* 0.1.

## Notes

### Competing Interest Statement

The authors have declared no competing interest.

